# Topology-based fluorescence image analysis for automated cell identification and segmentation

**DOI:** 10.1101/2022.06.22.497179

**Authors:** L. Panconi, M. Makarova, E. R. Lambert, R.C. May, D.M. Owen

## Abstract

Cell segmentation refers to the body of techniques used to identify cells in images and extract biologically relevant information from them; however, manual segmentation is laborious and subjective. We present Topological Boundary Line Estimation using Recurrence Of Neighbouring Emissions (TOBLERONE), a topological image analysis tool which identifies persistent homological image features as opposed to the geometric analysis commonly employed. We demonstrate that topological data analysis can provide accurate segmentation of arbitrarily-shaped cells, offering a means for automatic and objective data extraction. One cellular feature of particular interest in biology is the plasma membrane, which has been shown to present varying degrees of lipid packing, or membrane order, depending on the function and morphology of the cell type. With the use of environmentally-sensitive dyes, images derived from confocal microscopy can be used to quantify the degree of membrane order. We demonstrate that TOBLERONE is capable of automating this task.

## Introduction

Fluorescence microscopy is fundamental to modern cell biology [1]. However, in order to extract biologically relevant information from acquired images, researchers are required to first identify the regions of the image corresponding to individual cells or their enclosed organelles. The introduction of cell segmentation techniques allows for automated partitioning of images derived from fluorescence microscopy. In general, these techniques can be classified into machine learning-based methods and non-machine learning-based methods. Both variants present a range of drawbacks.

Non-machine learning methods typically require images with individual components of the cell stained specifically so that the algorithm can detect them [2]. They are usually dependent on cell geometry (in particular, convexity), making them incapable of identifying cells or organelles with particularly complex structures [3]. Additionally, they often require additional steps such as image reconstruction or seed-point extraction and typically have low generalisability for new applications [4, 5].

Machine learning-based methods employ a range of data manipulation techniques which learn from training data to make informed predictions on new images. In image analysis, these techniques are typically limited to supervised learning and therefore inherently require training data, using large sets of existing images which have already undergone cell segmentation [6]. Training data must typically be derived manually, which is a time-consuming process and creates subjectivity in the results. At the extremity of supervised object classification is deep learning. These methods carry several disadvantages in that they require training data to return accurate results and have a range of possible input parameters and settings (e.g. network architecture and training hyperparameters) [1, 7, 8]. Even with the advantages they bring, machine learning methods may also be limited to particular cell geometries [9].

In this work, we devise an image analysis algorithm for efficient cell and organelle segmentation irrespective of morphological and geometric distinctions. TOpological Boundary Line Estimation using Recurrence Of Neighbouring Emissions (TOBLERONE) enables identification of intensity modes within images by means of the topological data analysis techniques known as persistent homology and mode seeking. Topological image analysis is itself a relatively young field, which has shown promise in probing microbiology [10]. Similar algorithms have used this form of analysis to segment nuclei in histological slides of liver tissue [11]. Here, we have adapted this concept to identify any biological structure in fluorescence images.

Persistent homology was initially developed to study qualitative features of data sets [12], and then as a pre-requisite for cluster analysis, with filtrations constructed over a point cloud in which each localisation is assigned a local density given by the number of neighbouring points in a specified search radius [13]. In this instance, filtrations are constructed over a field of pixels with the “local density” derived exclusively from the fluorescence intensity of the underlying image.

To demonstrate the method on experimental, biological data we show that TOBLERONE can segment pixels corresponding to the plasma membrane of pan-membrane-stained cells. There is evidence that plasma membranes can comprise an extensive range of lipid packing states with varying levels of molecular order [14, 15]. It has been shown that such lipid packing states may be specifically regulated by cells during active cellular processes, suggesting that lipid-mediated membrane organisation has functional consequences [16, 17]. Using environmentally-sensitive membrane probes, we highlight the efficacy of TOBLERONE in mapping heterogeneity of lipid packing across the plasma membrane of the pathogenic yeast species, *Cryptococcus gattii*, and in mammalian cells.

## Materials and Methods

HEK293 cells were cultured in DMEM at 37°C in a 5% CO_2_ incubator. Cells were split and seeded into an 8-well coverslip bottomed Ibidi microscope chamber 24h prior to imaging. They were then stained with either 5μM di-4-ANEPPDHQ from a 5mM ethanol stock solution (to stain membranes) or 1X Nucblue (to stain DNA) 30mins prior to imaging. *Cryptococcus gattii* cells were cultured in yeast-peptone-dextrose (YPD) broth at 25°C with rotation and stained with di-4-ANEPPDHQ in the same way. Treatment with a hydroxyoleic acid (2μM) and 7-ketocholesterol (2μM) was performed 3 hours before imaging and staining.

Live HEK293 and *Cryptococcus gattii* cells were imaged on a Zeiss 780 laser-scanning confocal microscope at 37°C. For di-4-ANEPPDHQ, 488nm excitation was used with fluorescence collected in two wavelength ranges: 500-580nm and 620-750nm. GP values from these two images were calculated as previously described [15]. For Nucblue, excitation was at 405nm with fluorescence collected in the range 420-500nm. In both cases, 4X line averaging was used.

Simulated images were generated from arbitrary shapes in GIMP raster graphics editor. Poisson noise and Gaussian blur were applied using Fiji image manipulation software. The TOBLERONE software package was written in the R programming language, version 4.2.0, and employed in the integrated development environment RStudio. The code incorporated the tiff library for image manipulation and the built-in grid package for display. See Supplementary Material for code.

## Results

### Description of the algorithm

TOBLERONE is initialised with two inputs: the matrix representation of a grayscale image, where each pixel represents an intensity value between 0 and 1, and a parameter known as the *persistence threshold*, which dictates the permitted range of differences in intensity between pixels corresponding to the same object. Given that the image size and range of possible intensity values is finite, it is possible to produce a sequence of binarized images by thresholding over all possible intensities (Figure 1a-d). In order to undertake topological data analysis, each of these binary images must be ascribed an algebraic construct known as a simplicial complex. This is typically depicted in 2D space by a set of nodes (0-cells), connected by edges (1-cells) and spaces between three adjacent edges filled by a face (2-cells), giving the complex the appearance of a triangulation. To achieve this, TOBLERONE first maps each active pixel onto a node (or vertex) and assigns two nodes to be connected by an edge if they are immediately adjacent and are both active in the binary image – this representation is often referred to as the grid topology. The implementation of the grid topology (as the primary representation of an image) underpins the assumption that continuous objects are defined by a set of pixels in which a path between any two pixels can be achieved by a finite series of lateral or diagonal movements across pixels within the set.

**Figure 1:**
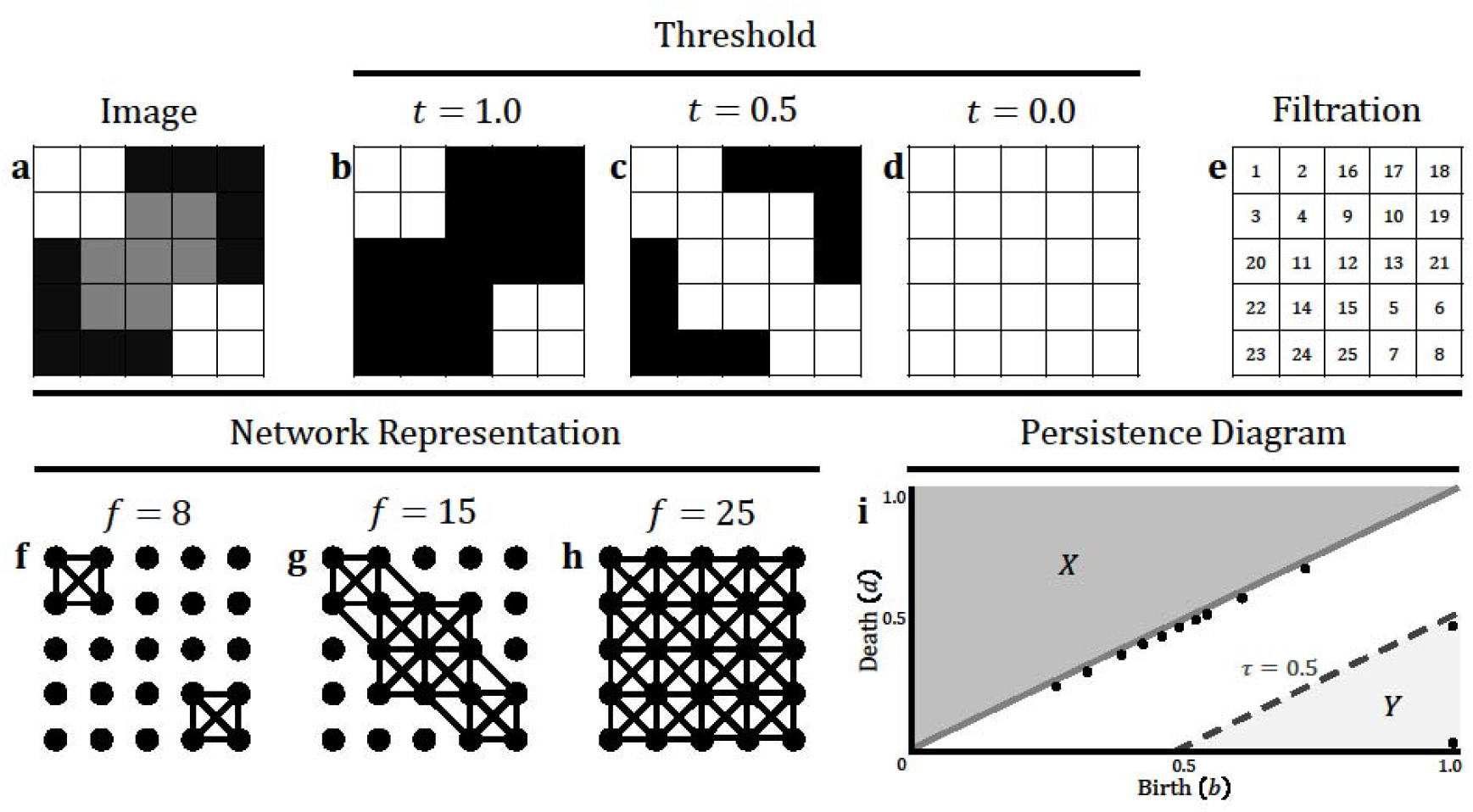
**a** A schematic image with 25 pixels, represented by a matrix in which each entry contains a numeric value between 0 and 1 corresponding to intensity. **b-d** Binary image at a threshold of *t* = 1.0, 0.5 and 0.0 respectively. **e** The filtrations matrix constructed by assigning each pixel a value corresponding to the order at which it was activated. Pixels which became active at the same threshold are numbered arbitrarily. **f-h** The corresponding network representation of the image at filtration values corresponding to the thresholds above. Here, *f* denotes the maximum filtration value permitted. As the filtration value increases, two connected components form and then merge into one object. **i** The persistence diagram constructed from the topological decomposition of the image. This plots the birth threshold, b, against the death threshold, *d*, for each object identified. The persistence of each object is represented by *τ* = *b* − *d*. Two prominent points are found in region *Y*, given by τ ≥ 0.5, which correspond to the two bright objects in the original image.

As we progress sequentially through the series of binary images (Figure 1b-d), the number of nodes, edges and faces within the graphical representation of the simplicial complex increases. As such, we can assign each cell complex an ordered numerical value corresponding to the stage of the binarization sequence at which it was first activated (Figure 1e). Mathematically, this is known as a filtration scheme, and each stage of the filtration will correspond to a different simplicial complex (Figure 1f-h) [18, 19]. The topological features of a simplicial complex can be described by the mathematical constructs known as homology classes. The topology of a given object can be described almost entirely by the homology classes associated with it [20, 21]. While the definition of a topological feature typically incorporates holes in the complex, we restrict our usage here to the first homology class, which represents the number of distinct connected components, and therefore omit 2-cells from our analysis [22]. Since the activation of an edge is imposed directly by the activation of the two nodes it is formed between, it suffices to observe the activation of pixels alone. As a result, we need only consider the filtration values of the nodes as their corresponding pixels are activated. Here, it is assumed that each identified connected component corresponds to a distinct biological structure.

By considering the image as a discretised representation of a continuous intensity field, we can identify the most prominent (i.e. brightest) pixels as estimations of the local maxima within that field. These maxima are known as modes, and are typically among the first nodes to appear within the filtration. As the filtration progresses, more nodes will be activated within the vicinity of the original mode and become connected. Since the mode is inherently defined as a local maximum, each newly added node will have a filtration value higher than that of the original. This induces a discretised gradient vector field upon the simplicial complex corresponding to the differences in filtration values between neighbouring nodes. To each mode we can then assign a *region of influence* corresponding to the set of nodes which are directed down the gradient field to the given mode. This mode is known as the *root* of the region of influence, with each root acting as the point of origin or *generator* for a connected component [13]. Neighbouring regions of influence are merged to form a single connected component with the new generator defined to be the youngest of the two original generators, that is, the root with the lowest filtration value. This process is known as mode-seeking and it is the principle which allows each connected component to be defined by a generator and its region of influence. This, in essence, encapsulates the mathematical notions which underpin the first homology class and allow for easy identification of connected components based on variations in pixel intensity.

At the end of the filtration, only one generator will remain. This corresponds to the pixel with highest intensity value, and its region of influence will incorporate all pixels in the image. This means that, throughout the process of generating the filtration, each root is formed and then subsequently destroyed, save for the final mode which will not be a good representation of the segmentation. As such, each root has an associated birth scale, the threshold at which the root and its corresponding connected component is created, and death scale, the threshold at which the region of interest intersects another and is absorbed. The difference between the birth and death scale is known as the *persistence*. This can be recorded for each connected component identified throughout the filtration and represented via a persistence diagram, which plots the birth scale against the death scale for each region [23]. Each point on this diagram corresponds to a connected component. We are particularly interested in those which have high birth scales and low death scales, as these correspond to regions of the space which form at high thresholds and do not merge with the background until particularly low thresholds are reached. By selecting an appropriate persistence threshold, we can extract only the most persistent connected components. A persistence diagram highlights the most persistent topological features of the image by plotting the birth scale against the death scale (Figure 1i). In a biological setting, this allows for identification and extraction of structures visible within the image, and essentially segments each object. After each implementation, TOBLERONE will return the number of connected components found with the given persistence threshold. An appropriate persistence threshold can be ascertained iteratively by arbitrarily initialising at τ = 0.5 and perturbing the threshold to match the expected number of connected components (in this context, the expected number of cells). Increasing the threshold will yield greater merging and reduce the number of connected components found, while decreasing the threshold will promote more pronounced segmentation and increase the number of connected components found.

Once the segmentation has been finalised, the algorithm will extract the exterior boundary of each object. However, probing micro-heterogeneity in membrane properties requires an additional understanding of the orientation of the circuit of pixels corresponding to the plasma membrane. As such, TOBLERONE employs a variation of the Swinging Arm method which preserves the ordering of each pixel comprising the boundary layer, initialised at the top-left-most pixel and proceeding counter-clockwise until returning to the origin [24]. This boundary can then be deleted from the image and the process can be repeated an arbitrary number of times, allowing the choice of layer selection at varying depths within the object. The algorithmic process culminates in the output of a given cell segmentation and the oriented loops corresponding to the boundaries of each found object.

### Demonstration with simulated data

Using a series of simulated images, with highly varied geometries and topologies, we have found that TOBLERONE can identify and segment binary images with high sensitivity and specificity (Figure 2a-f). A binary object was generated, and Poisson noise was simulated over each image with rate parameter *λ* = 1. Then, Gaussian blur was applied over a radius of *r* ≈ 14.14. These were chosen to approximately recapitulate the image quality obtained from conventional confocal microscopy. We then quantified TOBLERONE’s sensitivity (fraction of ground-truth bright pixels correctly labelled as active by the analysis) and the specificity (fraction of ground-truth dark pixels correctly analysed as off).

**Figure 2:**
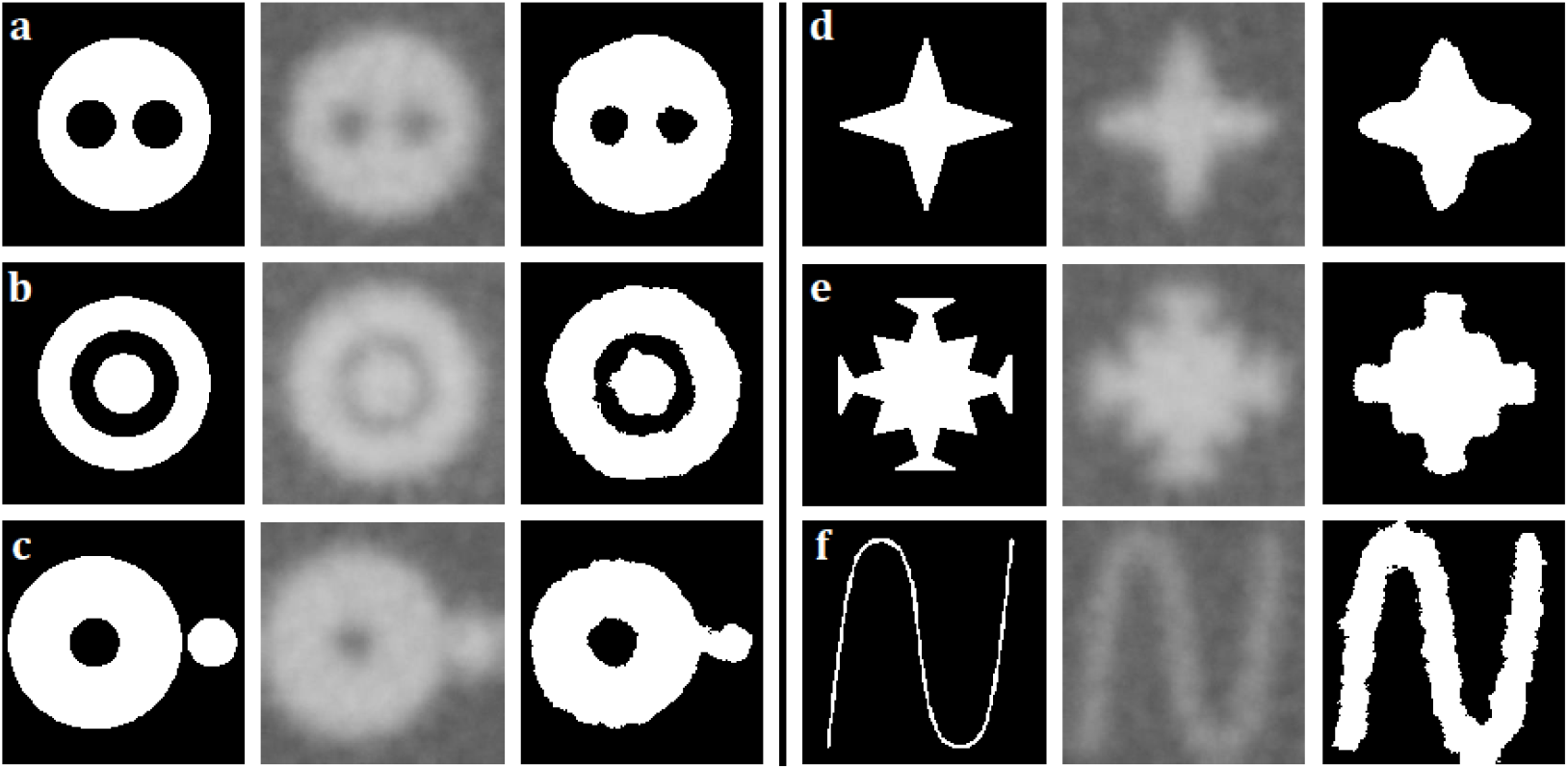
**a-f** A series of toy images exhibiting a range of geometries and topologies. Noise and blur are simulated over each image and then TOBLERONE is performed to recover the original segmentation. Primary segmentations were identified using a persistence threshold of 0.5.

Under these conditions, TOBLERONE achieved a sensitivity of 0.98 and a specificity of 0.87 (averaged over the 6 simulated image conditions) – this means that almost all pixels belonging to an object were correctly identified and most pixels belonging to the background were correctly ignored, respectively. In most instances, the algorithm successfully identified the correct number of individual connected components. This suggests that TOBLERONE is capable of accurately and precisely segmenting objects, even under poor image quality.

### Demonstration with experimental data

We then demonstrated TOBLERONE by using it to assess two cell types: the R265 strain of *Cryptococcus gattii*, which typically exhibits an approximately spherical geometry, and human embryonic kidney (HEK) cells, which display a more complex morphology incorporating finger-like protrusions. Segmentation results suggest that both HEK cells (Figure 3a-b) and their nuclei (Figure 3c-d) can be identified using TOBLERONE.

**Figure 3:**
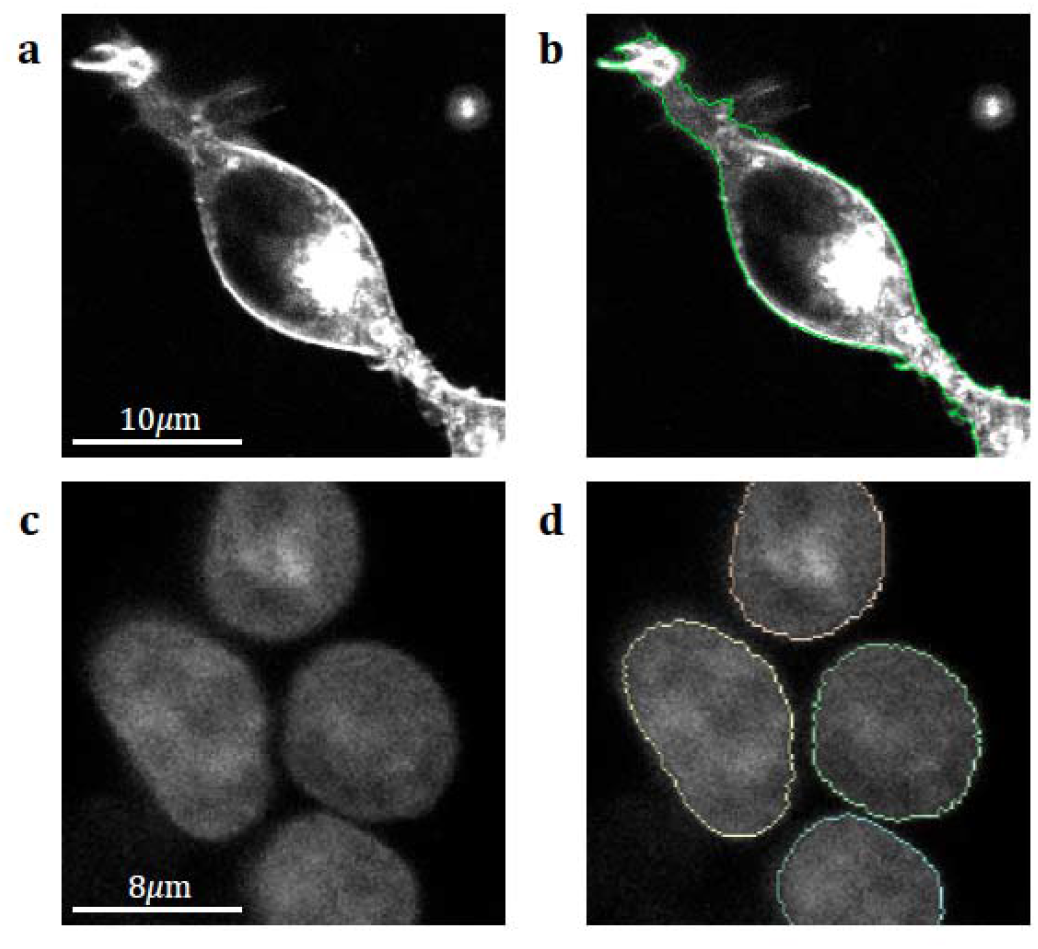
**a** Image of a HEK293 cell stained with di-4-ANEPPDHQ. **b** The boundary of the HEK cell as identified by TOBLERONE, overlaid onto the original image. Protrusions and variations in membrane morphology are captured despite the heterogeneous geometry. **c** An image of several HEK cell nuclei stained with Nucblue. **d** The boundaries of each nucleus as identified by TOBLERONE. Here, each boundary is highlighted as a separate connected component.

*C. gattii* is an infectious species of fungus which is responsible for cryptococcal meningitis in humans. TOBLERONE was used to identify the membranes of these cells (Figure 4a). The degree of membrane order, represented by the Generalised Polarisation (GP) image (Figure 4b), was calculated. The GP line profiles were then extracted from the boundaries (Figure 4c). The average GP was extracted for control cells as well as those treated with 2-hydroxyoleic acid or 7-ketocholesterol, which are predicted to introduce a higher degree of membrane disorder (Figure 4d). A statistically significant difference in the mean GP value was identified at the significance level of 1%, suggesting that both 2OHOA and 7-ketosterol reduce membrane order in *C. gattii*.

**Figure 4:**
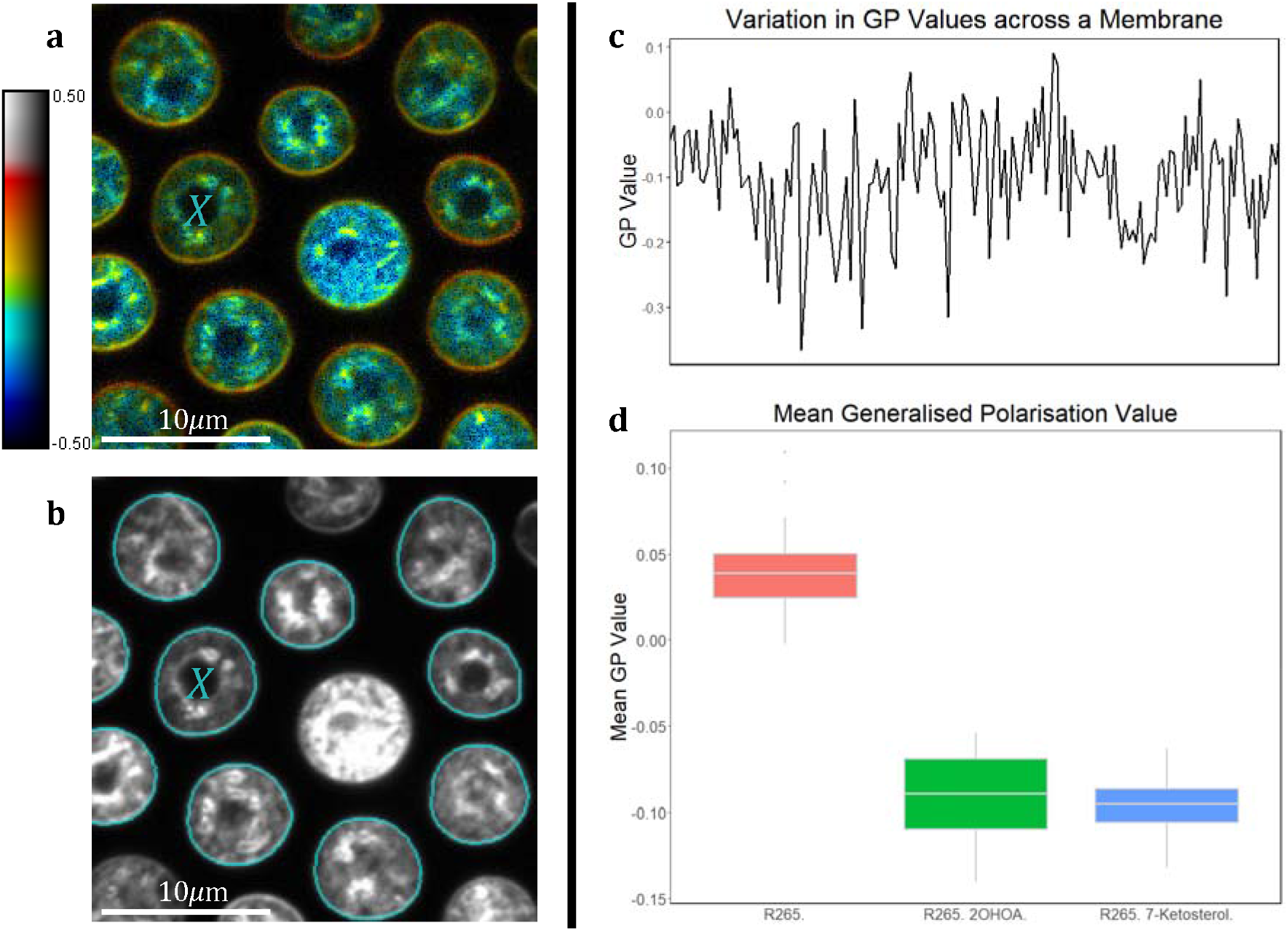
**a** A GP image of *C. gattii* cells stained with di-4-ANEPPDHQ. Psuedocolour applied to reflect the difference in GP values across the membrane. **b** The boundaries of the same cells, as found by TOBLERONE, overlaid onto the grayscale image in blue. Dead and incomplete cells lying on the periphery have been manually excluded. **c** The GP line profile extracted from the boundary of the cell marked *X*. **d** Boxplot depicting the average GP value across the membrane for cells treated with 2OHOA or 7-ketocholesterol.

## Discussion

In this work, we introduce TOBLERONE, an image segmentation algorithm specifically designed for identifying cells and organelles in fluorescence microscopy images, which operates without the drawbacks of conventional geometric and machine learning-based image segmentation methods. We have explored the mathematical principles which underpin the functionality of TOBLERONE, namely, in the use of topological data analysis for extracting the homology classes of the image under different thresholds. We then applied the algorithm to a range of simulated images – achieving high sensitivity and specificity even under particularly poor image quality – and a selection of differing cell and organelle geometries, including variants of the R265 strain of *C. gattii*, as well as human embryonic kidney cells and their nuclei. This demonstrated the applicability of TOBLERONE in practice by objectively identifying structures, biological or otherwise, of arbitrary shape. By exploiting topological properties of the boundaries of these structures, we were able to identify the plasma membrane of *C. gattii* and produce quantitative analysis of membrane order.

In comparison to the existing methods for image segmentation, TOBLERONE does present the following drawbacks. It is currently limited to 2D images, and its performance will be dependent on image quality [25]. Computationally, the primary segmentation’s runtime is on-par with existing machine learning methods, but pre-processing and boundary identification often take longer [6, 9]. While TOBLERONE can function with only a single parameter, it is not always obvious how to appropriately select its value – however, topological features in images are typically robust to variations in intensity, so a range of persistence thresholds will usually return a suitable segmentation [26-28]. As a topological image analysis technique, TOBLERONE is invariant of cell and organelle morphology [29, 30]. While this makes the algorithm highly generalisable, it can present challenges in adherent cell lines, as the algorithm may merge cells in close proximity, particularly when there is no visible gap between them.

The advantages of TOBLERONE over the general geometry-based approaches are as follows. TOBLERONE presents the advantage that it can segment structures of any size and shape, provided they are well-separated from surrounding objects and the background. In microscopy, this alleviates the need for pre-existing knowledge of the geometric properties of cells and, in the case of machine learning, training data sets of any kind. Not only does this counteract over-parameterisation, but it reduces the impact of parameter estimation. Additionally, since TOBLERONE is invariant of geometric cell properties, it is applicable even to images with a high degree of between-cell variation. As such, topological image analysis is best employed when the images contain structures of varied or complex geometry – particularly those with highly non-convex structures. Typically, it is more likely that geometric techniques would outperform topological methods in instances where cells form dense, connected tissues.

Generally, TOBLERONE probes images containing structures of any topology, including those which appear to present gaps within the structures themselves. This principle allows for the identification of both exterior boundaries, corresponding to the cell membrane, and interior boundaries, which may correspond to organelles, vesicles or other structures contained within the cells. As TOBLERONE is a topological methodology, it is able to distinguish between the different boundary types automatically. Overall, this tool presents a different approach to image analysis compared to geometrical or machine-learning based image segmentation with considerable advantages for identifying cell boundaries.

## Acknowledgments

We thank Dr. Daniel Nieves and Dr. Jeremy Pike for critically reading the manuscript. We acknowledge the Microscopy and Imaging Services at Birmingham University (MISBU) for microscope access. We acknowledge funding from Oxford Nanoimaging (ONI) and the Engineering and Physical Sciences Research Council via the University of Birmingham CDT in Topological Design.

## Supplementary Material

The TOBLERONE R package, with a given example dataset, is available at https://github.com/lucapanconi/toblerone.

